# Exploration of *Methanomethylophilus alvus* pyrrolysyl-tRNA synthetase activity in yeast

**DOI:** 10.1101/2022.01.07.475408

**Authors:** Jessica T. Stieglitz, Priyanka Lahiri, Matthew I. Stout, James A. Van Deventer

## Abstract

Archaeal pyrrolysyl-tRNA synthetases (PylRSs) have been used to genetically encode over 200 distinct noncanonical amino acids (ncAAs) in proteins in *E. coli* and mammalian cells. This vastly expands the range of chemical functionality accessible within proteins produced in these organisms. Despite these clear successes, explorations of PylRS function in yeast remains limited. In this work, we demonstrate that the *Methanomethylophilus alvus* PylRS (MaPylRS) and its cognate tRNA_CUA_ support the incorporation of ncAAs into proteins produced in *S. cerevisiae* using stop codon suppression methodologies. Additionally, we prepared three MaPylRS mutants originally engineered in *E. coli* and determined that all three were translationally active with one or more ncAAs, although with low efficiencies of ncAA incorporation in comparison to the parent MaPylRS. Alongside MaPylRS variants, we evaluated the translational activity of previously reported *Methanosarcina mazei*, *Methanosarcina barkeri*, and chimeric *M. mazei* and *M. barkeri* PylRSs. Using the yeast strain RJY100, and pairing these aaRSs with the *M. barkeri* tRNA_CUA_, we did not observe any detectable stop codon suppression activity under the same conditions that produced moderately efficient ncAA incorporation with MaPylRS. The addition of MaPylRS to the orthogonal translation machinery toolkit in yeast potentially opens the door to hundreds of ncAAs that have not previously been genetically encodable using other aminoacyl-tRNA synthetase/tRNA pairs. Extending the scope of ncAA incorporation in yeast could powerfully advance chemical and biological research for applications ranging from basic biological discovery to enzyme engineering and therapeutic protein lead discovery.

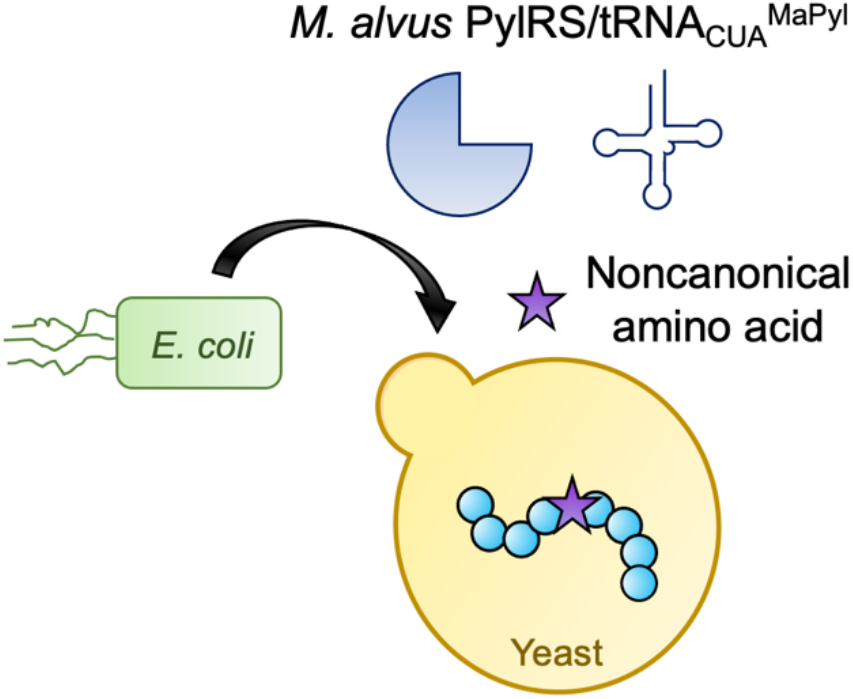

## Introduction

Several archaeal pyrrolysyl-tRNA synthetases (PylRSs) facilitate expansion of the genetic code of *E. coli* and mammalian cells, allowing for the site-specific incorporation of hundreds of noncanonical amino acids (ncAAs) into proteins in response to stop codons.^*1–4*^ The ability to precisely control the location(s) of ncAAs with wide ranges of chemical functionality significantly advances applications in synthetic biology, chemical biology, and protein engineering. One of the major advantages of using archaeal PylRS and tRNA^Pyl^ pairs is that they do not cross-react with any of the endogenous aminoacyl-tRNA synthetases (aaRSs) or tRNAs in *Escherichia coli*, yeast, or mammalian cells. This orthogonality to the protein translation machinery in non-archaeal organisms has supported PylRS engineering in *E. coli* and transfer of *E. coli*-engineered PylRSs to mammalian cells.^*5*^ Unlike the other commonly utilized aaRSs in *E. coli* and eukaryotes, wild-type PylRSs exhibit high levels of substrate polyspecificity.^*2*^ The plasticity of PylRS active sites have already supported the genetic encoding of a depth and breadth of ncAA functionalities that are not currently accessible using alternative aaRSs in either prokaryotes or eukaryotes.

The most commonly employed PylRSs are the *Methanosarcina mazei* and *Methanosarcina barkeri* PylRSs (MmPylRS and MbPylRS). MmPylRSs and MbPylRSs have been engineered in *E. coli* and mammalian cells to vastly expand the collection of ncAAs that can be genetically encoded in proteins.^*6*–*10*^ The ability to evolve PylRSs in *E. coli* and then transfer the orthogonal translation systems (OTSs) directly to mammalian cells bypasses the need to engineer such aaRSs directly in mammalian cell lines. However, there are some notable challenges to working with PylRSs, such as their insoluble N-terminal regions,^*11*, *12*^ which can lead to insolubility even following codon optimization in *E. coli.^13^* On the other hand, yeast such as *Saccharomyces cerevisiae* fall into a unique category between *E. coli* and mammalian cells. Similar to *E. coli, S. cerevisiae* exhibits rapid doubling times during growth and is well-characterized as a model organism. However, unlike *E. coli*, yeast can efficiently produce complex proteins such as antibodies and provide access to eukaryotic biology that is conserved or at least similar to the biology of mammalian cells. Furthermore, powerful tools and methods exist in yeast such as 1) yeast knockout collections and other genetic resources that facilitate deep understanding of biological phenomena; and 2) yeast display and other engineering tools that facilitate protein engineering, metabolic engineering, and synthetic biology.^*14*–*19*^

There are few reports evaluating MmPylRS and MbPylRS activities in yeast. Yokoyama and coworkers utilized two mutant MmPylRSs to encode Boc-L-lysine and *N*_ε_-benzoyl-L-lysine (BocK and LysZ, respectively), in *S. cerevisiae* MaV203 using a tRNA^Val^-tRNA^MmPyl^ system.^*8*^ With 1 mM ncAA concentrations during induction of protein synthesis, low levels of TAG codon suppression were detected. Later, Chin and coworkers reported activity of multiple variants including MbPylRS/tRNA^MbPyl^ and MbPylRS/tRNA^MmPyl^ pairs in *S. cerevisiae* MaV203:pGADGAL4(2TAG).^*20*^ Using MbPylRS mutants, the researchers were able to demonstrate incorporation of five ncAAs using concentrations during induction of protein synthesis ranging from 1.3–10 mM. One additional report by Kapoor and coworkers in 2013 utilized MbPylRSs^*20*, *21*^ to encode three ncAAs using 2–10 mM concentrations during induction of protein expression in *S. cerevisiae* INVSc1, with low but detectable incorporation.^*22*^ Unlike in *E. coli* and higher order eukaryotes, use of MmPylRSs and MbPylRSs in yeast appears to require higher ncAA concentrations^*20*, *23*^ while yielding low to moderate levels of ncAA incorporation. These findings motivate the exploration of PylRSs from other organisms in yeast in search of OTSs that exhibit ncAA incorporation efficiency and translational activity with a broad range of ncAAs.

Recently, Chin and coworkers identified several methanogen PylRSs capable of supporting stop codon readthrough with the ncAA Boc-lysine (BocK) that exhibited orthogonality in *E. coli* and did not mischarge cAAs.^*23*^ One particularly active PylRS was from *Methanomethylophilus alvus* (MaPylRS), and the MaPylRS/tRNA_CUA_^MaPyl^ pair has since been demonstrated to be highly translationally active in both *E. coli* and mammalian cells.^*24*–*27*^ MaPylRS lacks the insoluble N-terminal domain that was previously considered to be essential for tRNA recognition^*11*, *28*^ and shares a homologous active site with PylRSs known to be active in other organisms. This suggests that it may be a suitable starting point for evolving variants that accept a similarly broad range of ncAAs known to be substrates of engineered PylRS variants from *M. barkeri* and *M. mazei*. Despite the high translational activity of MaPylRS in *E. coli* and mammalian systems, its activity has not been evaluated previously in yeast to the best of our knowledge.

Here, we evaluate the translational activity of MaPylRS in *S. cerevisiae* alongside several *M. mazei* and *M. barkeri* PylRSs. We first determined that wild-type MaPylRS supports protein translation with several ncAAs with varying levels of stop codon readthrough efficiency in yeast. Given the availability of numerous previously reported variants of MaPylRS with known translational activity in *E. coli* and mammalian cells, we investigated the activities and substrate preferences of these variants in yeast. All three variants we tested exhibited low but detectable levels of translational activity with one or more ncAAs. In contrast, none of the MmPylRS or MbPylRS variants tested under the same conditions as MaPylRS variants in this work showed detectable translational activity. The availability of translationally active MaPylRS variants in yeast provides potential access to vast sets of ncAAs that have not previously been genetically encoded in this organism. Given the importance of yeast in basic biology and as a chassis for protein engineering and synthetic biology, a larger chemical toolkit is expected to support advances ranging from a better understanding of eukaryotic biology to the identification of new therapeutic targets and discovery of new therapeutic leads. Access to additional active PylRSs in yeast drastically expands the ncAA incorporation landscape in this organism and enables numerous applications that rely on the efficient genetic incorporation of ncAAs.

## Results and Discussion

### Evaluating the activity of *M. alvus* PylRS in yeast

We first sought to determine whether the wild-type MaPylRS is active in *S. cerevisiae* RJY100, a commonly used yeast strain.^*29*^ MaPylRS and tRNA_CUA_^MaPyl^ were expressed under constitutive GPD and SNR52 promoters, respectively, alongside a galactose-inducible dual-fluorescent protein reporter system for evaluating ncAA incorporation (Fig. 1A and B). The reporter consists of blue fluorescent protein (BFP) fused to GFP by a flexible linker sequence that either contains (BXG) or does not contain (BYG) a TAG codon.^*30*^ Dual-terminus detection reporting platforms allow for the evaluation of ncAA incorporation efficiency and fidelity using the metrics relative readthrough efficiency (RRE) and maximum misincorporation efficiency (MMF).^*30*–*32*^ RRE is a metric of how efficiently a stop codon is read through in comparison to wild-type protein translation, while MMF is a metric to evaluate worst-case canonical amino acid incorporation levels in the absence of ncAAs.

**Figure 1.**
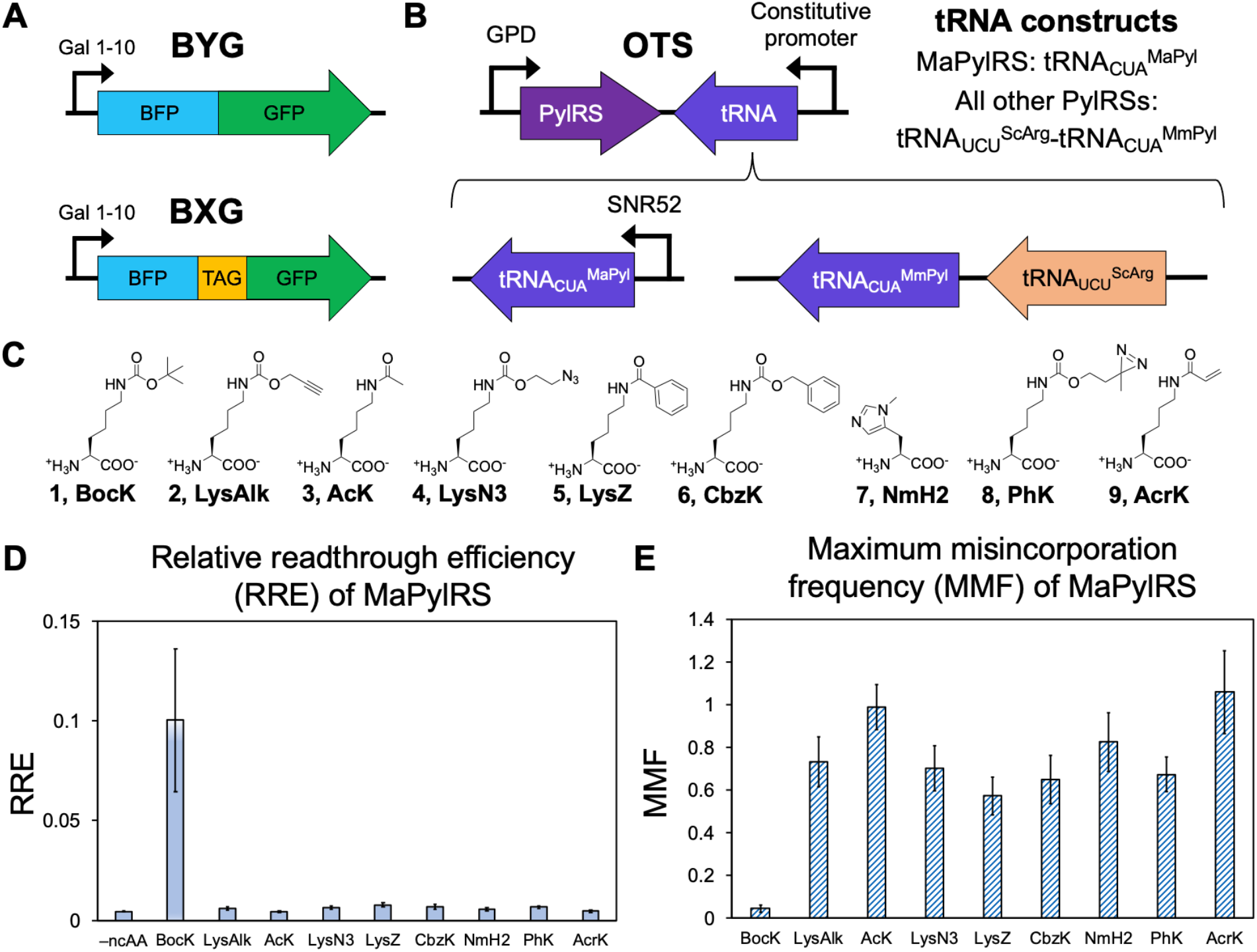
Efficiency and fidelity of wild-type MaPylRS with nine ncAAs. A) Schematic diagrams of BFP-GFP reporter constructs. BYG expresses BFP linked to GFP with no TAG codon under a galactose-inducible promoter. BXG is identical to BYG except that it contains a TAG codon at the last position of the linker sequence. B) The orthogonal translation system (OTS) plasmid expresses the PylRS and tRNA under constitutive promoters. For MaPylRS, tRNA_CUA_^MaPyl^ was co-expressed under the SNR52 promoter. For all other *M. mazei*, *M. barkeri*, or chimeric *M. mazei* and *barkeri* PylRSs, a two-tRNA expression system was used. In this system, expression of *S. cerevisiae* tRNA_UCU_^Arg^ drives expression of *M. mazei* tRNA_CUA_^Pyl^.^*20*^ C) Structures of ncAAs. **1**: Boc-L-lysine (BocK); **2**: 2-Amino-6-(prop-2-ynoxycarbonylamino)hexanoic acid (LysAlk); **3**: *N*_ε_-acetyl-L-lysine (AcK); **4**: (*S*)-2-amino-6-((2-azidoethoxy)carbonylamino)hexanoic acid (LysN3); **5**: *N*_ε_-benzoyl-L-lysine (LysZ); **6**: *N*_ε_-benzyloxycarbonyl-L-lysine (CbzK); **7**: 3-methyl-L-histidine (NmH2); **8**: (*S*)-2-Amino-6-((2-(3-methyl-3H-diazirin-3-yl)ethoxy)carbonylamino)hexanoic acid (PhK); and **9**: acryloyl-L-lysine (AcrK). D) RRE of cells transformed with a plasmid encoding MaPylRS induced in the absence of ncAAs (–ncAA) and in the presence of 10 mM concentration of each of the indicated ncAAs. E) MMF of cells constitutively expressing MaPylRS for each of the indicated ncAAs.

Cells containing the MaPylRS and reporter constructs were induced in the presence of each of nine ncAAs supplemented into the media at a 10 mM final concentration: **1**: Boc-L-lysine (BocK); **2**: 2-Amino-6-(prop-2-ynoxycarbonylamino)hexanoic acid (LysAlk); **3**: *N*_ε_-acetyl-L-lysine (AcK); **4**: (*S*)-2-amino-6-((2-azidoethoxy)carbonylamino)hexanoic acid (LysN3); **5**: *N*_ε_-benzoyl-L-lysine (LysZ); **6**: *N*_ε_-benzyloxycarbonyl-L-lysine (CbzK); **7**: 3-methyl-L-histidine (NmH2); **8**: (*S*)-2-Amino-6-((2-(3-methyl-3H-diazirin-3-yl)ethoxy)carbonylamino)hexanoic acid (PhK); and **9**: acryloyl-L-lysine (AcrK) (Fig. 1C). Flow cytometry analysis of BFP and GFP fluorescence levels revealed substantial BocK incorporation, with an RRE value of 0.10 ± 0.036 and MMF value of 0.044 ± 0.016 (Fig. 1D and E, SI Figs. 1 and 2). Low levels of incorporation of LysAlk, LysN3, LysZ, and PhK were detected qualitatively in flow cytometry dot plots, but not at levels that shifted RRE values above the observed background levels (SI Fig. 2). To our knowledge, this is the first evidence of MaPylRS TAG codon suppression in yeast, and these promising findings prompted us to further investigate the potential utility of MaPylRSs in yeast.

### Comparing MaPylRS with MmPylRSs and MbPylRSs

After demonstrating that MaPylRS exhibits translational activity in yeast with several ncAAs, we compared the measured activities of MaPylRS against several *M. mazei* and *M. barkeri* PylRSs in RJY100. We chose the wildtype MbPylRS and MmPylRS as well as an additional two chimeric PylRSs^*33*^ (comprised of residues 1–149 of MbPylRS and 185–454 of MmPylRS): chPylRS and chAcK3RS. Previous reports demonstrated that MmPylRS^*23*^, MbPylRS,^*33*^ and chPylRS^*33*^ support translation with BocK, while chAcK3RS supports translation with AcK.^*33*^ Cells containing each of the five PylRSs and the corresponding tRNA construct (SNR52-tRNA_CUA_^MaPyl^ for MaPylRS; tRNA_UCU_^ScArg^-tRNA_CUA_^MmPyl^ for all other PylRSs) were induced in the presence of 10 mM BocK, AcK, and LysAlk and evaluated for GFP detection to identify successful readthrough events. GFP levels above background were detected when cells constitutively expressing MaPylRS were induced in the presence of BocK, indicating moderately high translational activity (Fig. 2, SI Fig. 3). Additionally, low but detectable levels of readthrough were observed when cells containing MaPylRS were induced in the presence of LysAlk but not AcK. In contrast, no detectable readthrough was observed in cells expressing any of the other four PylRSs following induction in the presence of BocK, AcK, or LysAlk (Fig. 2, SI Fig. 3). While our observations result in apparent discrepancies with previously reported findings related to PylRS activities in yeast,^*20*, *22*, *34*^ we note several differences between our experiments and the experimental conditions used in prior reports. In particular, changes in yeast strains, reporter system plasmid architecture, and orthogonal translation system plasmid architecture provide numerous potential explanations for the different experimental outcomes. Regardless, our demonstration of translational activity of MaPylRS in a common yeast display strain is notable because access to chemically diversified proteins in yeast display format is of potentially broad utility in protein engineering and chemical biology.

**Figure 2.**
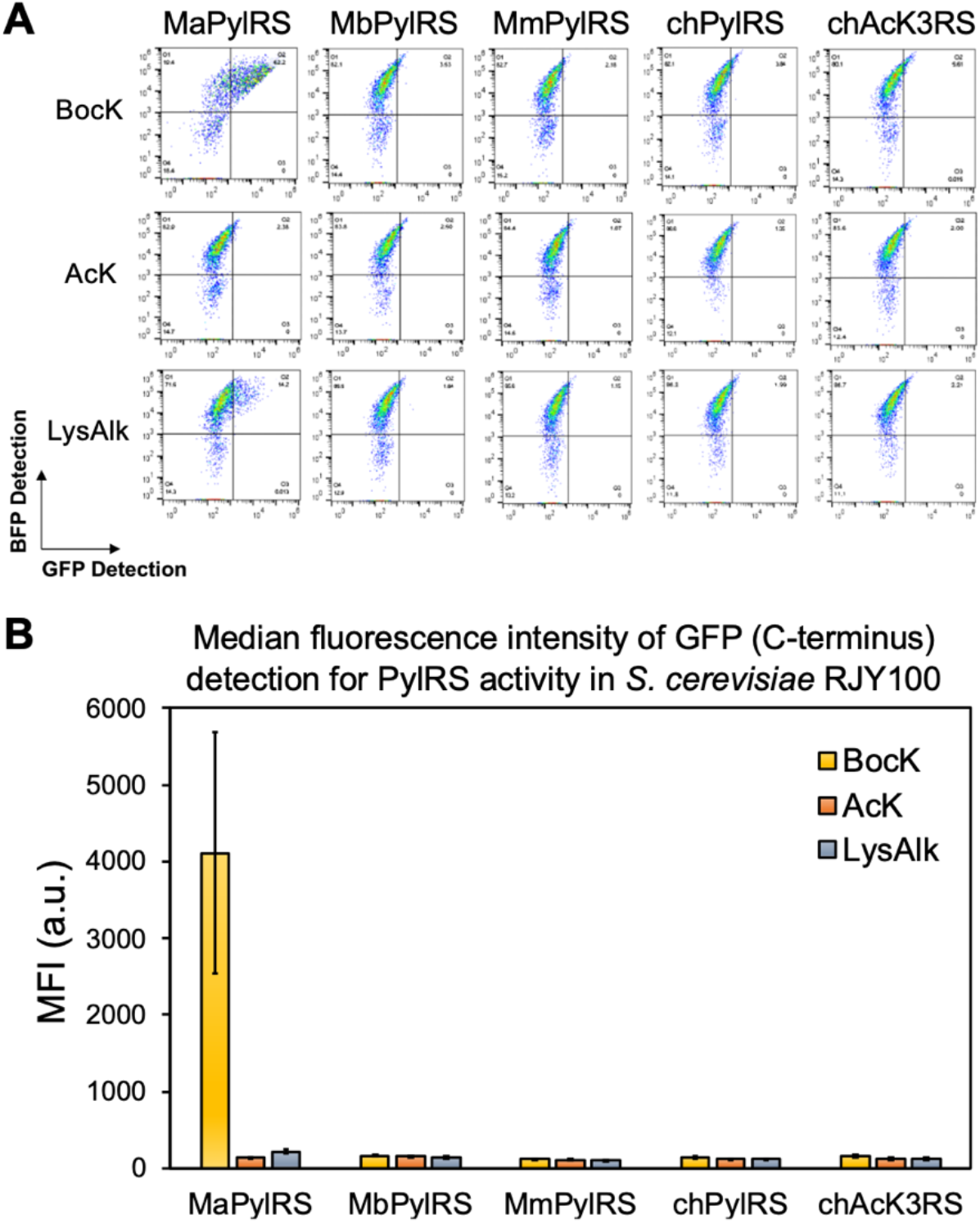
Comparison of activity of various archaeal PylRSs with cognate ncAAs. A) Flow cytometry dot plots for the yeast strain RJY100 transformed with plasmids encoding each of five PylRSs derived from *M. alvus*, *M. barkeri*, and *M. mazei* induced in the presence of 10 mM BocK, AcK, or LysAlk. B) Median fluorescence intensity (MFI) of GFP (full-length dual fluorescent reporter) detection for each condition in part A. For each population, the autofluorescence determined from uninduced cells was subtracted from the average of three biological replicates depicted here. Error bars represent the standard deviation of the biological triplicates. Control samples induced in the absence of ncAAs can be found in SI Figure 2.

### Testing the translational activity of three MaPylRS mutants

We next sought to determine if previously described MaPylRS variants used in other hosts would also exhibit translational activity in yeast. To investigate this question, we constructed three MaPylRS mutants originating from engineering efforts in *E. coli*^*23*, *24*^ or mammalian^*25*^ hosts (SI Table 1) and characterized their translational activity in yeast. MaPylRS-Mut1 contains mutations Y126M, M129G, and V168T and is known to encode CbzK;^*23*^ MaPylRS-Mut2 contains mutations L121M, L125I, Y126F, M129A, and V168F and encodes ncAA NmH2;^*24*^ and MaPylRS-Mut3 has a Y126A mutation and is active with axial *trans*-cyclooct-2-ene-L-lysine (TCO*K), *N*_ε_-[[(2-methyl-2-cyclopropene-1-yl)methoxy] carbonyl-L-lysine (CpK), exo-BCN-L-lysine (BCNK), and *N*_ε_-[(2-(3-methyl-3H-diazirin-3-yl)ethoxy)carbonyl]-L-lysine (AbK).^*25*, *35*, *36*^ We evaluated all three mutants with nine ncAAs added to induction media at 10 mM final concentrations (SI Fig. 4). Use of the dual-fluorescent reporter system described above identified MaPylRS-Mut1 as exhibiting detectable readthrough with CbzK; MaPylRS-Mut2 as exhibiting readthrough with NmH2 and AcrK; and MaPylRS-Mut3 as exhibiting readthrough with CbzK and PhK. We next determined the RRE and MMF of these combinations of mutants and ncAAs. These experiments revealed that while qualitative readthrough is apparent in flow cytometry dot plots, calculated RRE values were all at or only slightly above controls, with MMF values exhibiting the correspondingly high values that typically accompany low RRE values (Fig. 3, SI Figs. 5 and 6).^*30*, *32*^ These observations are consistent with our prior work and unpublished studies,^*37*^ which indicate that data collected with aaRSs exhibiting low or variable readthrough activity yields RRE values similar to background levels. In any case, our evidence indicating MaPylRS translational activity with several ncAAs hints at the potential to expand access to genetically encodable functionalities in yeast, especially if the translational activities of these mutants can be improved in future work.

**Figure 3.**
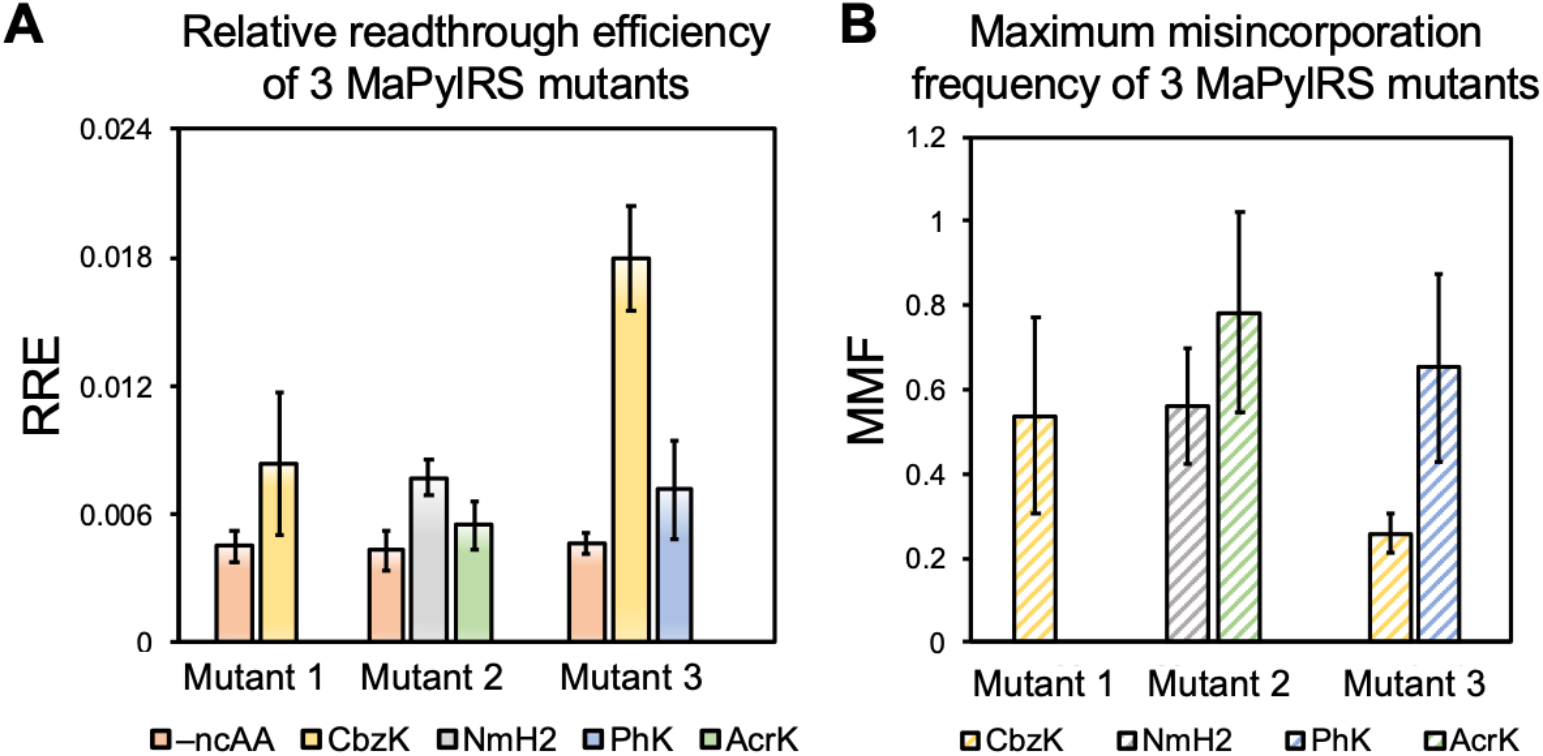
Efficiency and fidelity of ncAA incorporation in cells containing each of three MaPylRS mutants induced in the presence of 10 mM ncAA. A) RRE determined following induction of cells in the presence of 10 mM of each of the indicated ncAAs and absence of ncAAs (–ncAA). B) MMF determined with cells constitutively expressing each of the MaPylRS variants for the specified ncAAs.

### Mass spectrometry validation of MaPylRS activity in yeast

Finally, to validate incorporation of BocK by MaPylRS in yeast, we expressed and purified a protein reporter from RJY100 with a TAG codon at a permissible site. The reporter protein is an scFv-Fc expressed from a secretion vector, where the antibody variable fragment (scFv) is a Donkey IgG binder (Donkey1.1) that contains a TAG codon at the 54^th^ position in the heavy chain (Donkey1.1-H54TAG).^*38*^ 500 mL (Donkey1.1) or 1 L (Donkey1.1-H54TAG) cultures of RJY100 were induced for four days in YPG media, a standard rich media for induction, to provide a stringent evaluation of the MaPylRS variants’ ability to encode the ncAA for which they demonstrated the highest level of activity from previous experiments outlined above. Wild-type Donkey1.1 was induced in the absence of ncAAs; all Donkey1.1-H54TAG reporters expressed in cells containing MaPylRS variants were induced in the presence of 10 mM ncAA. Secreted proteins were harvested and purified via Protein-A column chromatography. An attempt was made to purify the Donkey1.1-H54TAG reporter protein with MaPylRS-Mut2 induced in the presence of 10 mM NmH2 and MaPylRS-Mut3 induced in the presence of 10 mM CbzK, but the isolated protein yield was too low to proceed with mass spectrometry analysis. However, wild-type Donkey1.1 and Donkey1.1-H54TAG expressed from cells containing MaPylRS and induced in the presence of 10 mM BocK were successfully expressed (SI Fig. 7).

The wild-type and H54TAG reporter proteins were concentrated and then enzymatically digested using trypsin prior to evaluation via matrix-assisted laser desorption ionization (MALDI) mass spectrometry (Fig. 4). For the desired site for ncAA incorporation, the expected peptide mass was 2234.1 Da for the wildtype sequence and the observed mass was at 2234.2 Da. The peak at 2211.3 Da in the WT plot is due to trypsin autolysis.^*39*^ The expected mass for the peptide with BocK encoded at the H54TAG codon was 2375.3 Da with an observed mass of 2375.8 Da. The expected peptides were present for both the wild-type and BocK-containing peptides, indicating that MaPylRS successfully charged BocK for incorporation at the TAG codon of interest. Although future work will be needed to improve production of soluble forms of ncAA-containing proteins with PylRSs in yeast, the data presented in this study confirm that MaPylRS is sufficiently active in *S. cerevisiae* RJY100 to enable production of a BocKcontaining protein, expanding the collection of aaRSs that can be utilized for genetic code expansion in yeast.

**Figure 4.**
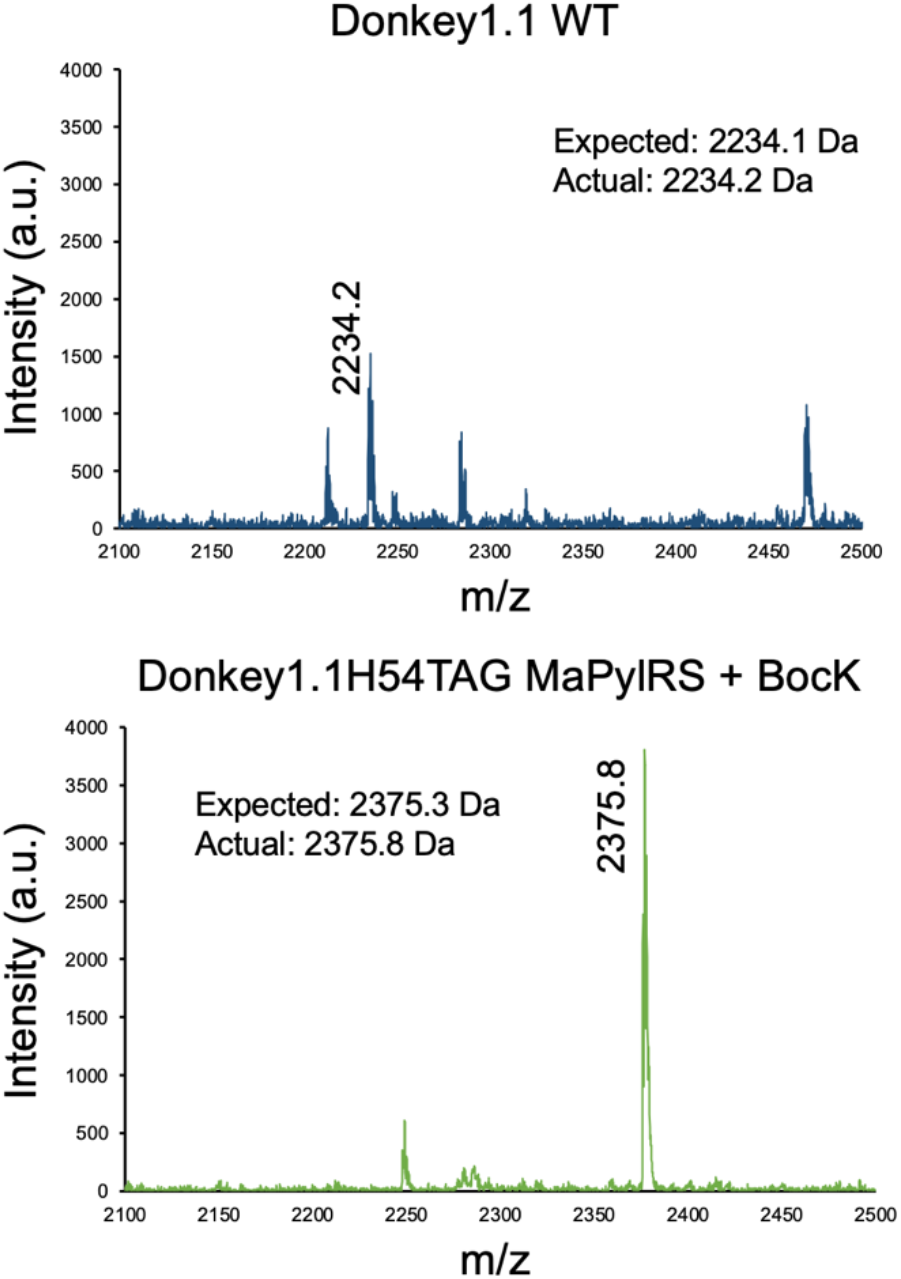
MALDI mass spectrometry spectra of a tryptic-digested WT reporter protein (Donkey1.1) and TAG codon-containing reporter (Donkey1.1-H54TAG) induced with MaPylRS and 10 mM BocK.

## Conclusions

For the first time, we present evidence that MaPylRS/tRNA_CUA_^MaPyl^ exhibits translational activity with several ncAAs in *S. cerevisiae*. Use of a dual-fluorescent reporter and the yeast display strain RJY100 enabled identification of MaPylRS activity with five of the ncAAs tested in this work. Incorporation of BocK, which occurred at the highest measured efficiency, was confirmed via MALDI mass spectrometry. Additionally, three MaPylRS mutants exhibited translational activity with various ncAAs. Many of these activities had been previously identified in organisms other than yeast, but at least two of the ncAAs evaluated here had not previously been reported to be incorporated into proteins produced in yeast: AcrK and NmH2. Using the same yeast strain (RJY100), ncAAs (BocK, AcK, and LysAlk), dual-fluorescent reporter, and previously described MmPylRS, MbPylRS, and chimeric MmPylRS/MbPylRSs, we were not able to detect ncAA incorporation activity, even for PylRSs that have shown activity with the same ncAAs in other yeast strains. Our findings here are a critical step toward improving the accessibility of PylRSs for use in yeast to facilitate incorporation of diverse ncAAs that have only been accessible in *E. coli* and mammalian systems.

Archaeal PylRSs have proved invaluable for site-specifically encoding unique and functionally advantageous ncAAs to advance applications that use *E. coli* or higher order eukaryotic platforms for protein expression. In yeast, PylRSs are relatively underdeveloped, with only a few reports of PylRS activity in the *S. cerevisiae* strains MaV203 and INVSc1. The activity of MaPylRS and variants in RJY100 is lower than previously reported *E. coli* aaRSs, despite the use of ncAA concentrations ten times higher than the concentrations generally used with *E. coli* aaRSs in yeast. In addition, we note that the highest levels of activity observed with the combinations of variants and ncAAs investigated here were substantially lower than the activity of the wild-type MaPylRS with BocK. This suggests that MaPylRS variants may need to be evolved in yeast in order to improve their activities, especially with regarding to required ncAA concentrations during induction. Auspiciously, *S. cerevisiae* is a robust organism for engineering enzymes and other binding proteins, with potent high-throughput technologies such as fluorescence-activated cell sorting in combination with yeast display.^*37*, *38*^ In addition, Chin and coworkers recently reported a substantial number of additional PylRSs that exhibit translational activity in *E. coli.^23^* Evaluating the performance of these aaRSs in yeast may lead to the identification of PylRSs that exhibit even higher levels of translational activities that what we observed in these studies. Exploration of the orthogonality of MaPylRSs with *E. coli* aaRSs in yeast could lead to systems supporting incorporation of multiple distinct ncAAs into proteins for applications including the examination of protein-protein interactions,^*40*–*42*^ among other potential uses. Expanding the limited set of distinct aaRSs available in yeast allows access to a broader set of genetically encodable ncAAs, which in turn is expected to impact many critical applications from epigenetics to biological therapeutic discovery.

## Materials and Methods

### Materials

Synthetic oligonucleotides for cloning and sequencing were purchased from GENEWIZ. Restriction enzymes used for vector digests were purchased from New England Biolabs (NEB). Sanger sequencing was performed by Quintara Biosciences (Cambridge, MA). Epoch Life Science GenCatch™ Plasmid DNA Mini-Prep Kits were used for plasmid DNA purification from *E. coli* and Zymo Research Frozen-EZ Yeast Transformation II kits were used to prepare yeast chemically competent cells and perform plasmid transformations. Noncanonical amino acids were purchased from the following companies: **1**: Boc-L-lysine (BocK, Chem-Impex International); **2**: 2-Amino-6-(prop-2-ynoxycarbonylamino)hexanoic acid (LysAlk, AstaTech); **3**: *N*__ε__-acetyl-L-lysine (AcK, Sigma-Aldrich); **4**: (*S*)-2-amino-6-((2-azidoethoxy)carbonylamino)hexanoic acid (LysN3, Iris Biotech GmBH); **5**: *N*_ε_-benzoyl-L-lysine (LysZ, Sigma-Aldrich); **6**: *N*_ε_-benzyloxycarbonyl-L-lysine (CbzK, Alfa Aesar™); **7**: 3-methyl-L-histidine (NmH2, Chem-Impex International); **8**: (*S*)-2-Amino-6-((2-(3-methyl-3H-diazirin-3-yl)ethoxy)carbonylamino)hexanoic acid (PhK, Iris Biotech GmBH); and **9**: acryloyl-L-lysine (AcrK, Iris Biotech GmBH).

### Media preparation and yeast strain construction

All experiments in yeast were performed in the *S. cerevisiae* strain RJY100, the construction of which has been described in detail elsewhere.^*29*^ Liquid and solid media were prepared as described previously.^*32*^ SD-SCAA and SG-SCAA media were prepared with the following: tryptophan (TRP), leucine (LEU), and uracil (URA) were omitted from the amino acid dropout mix for all cases in which cells were co-transformed with pRS315 (LEU2 marker) and pCTCON2 (TRP1 marker) or pCHA (TRP1 marker). SD-CAA (–TRP –ADE –URA) was used for propagation of pCHA-Donkey1.1 WT control for mass spectrometry experiments. 50 mM L-isomer ncAA stocks were prepared by adding DI water to the solid ncAA to 90% of the final volume, then utilizing 6.0 N NaOH as needed to dissolve the ncAA. DI water was added to the final volume and the stocks were sterile filtered through a 0.2 μm filter and stored at 4°C for up to two weeks prior to induction.

### Plasmid construction

The construction of vectors pCTCON2-BXG (Addgene plasmid #158127),^*30*^ pCTCON2-BYG (Addgene plasmid #158144),^*30*^ pCHA-Donkey1.1,^*38*^ and pCHA-Donkey1.1-H54TAG^*38*^ have been described previously. MmPylRS was cloned by inserting a codon-optimized gene block into pRS315-AcFRS. The MmPylRS gene block was PCR-amplified with primers containing 30 bp overlap with pRS315 double restriction enzyme-digested with NcoI and NdeI. Both the amplified MmPylRS gene and vector were analyzed on an agarose gel, extracted, and ligated together. The ligation was transformed into *E. coli*, plated on selective media (50 μg/mL ampicillin), and grown overnight. Colonies from the selective media plate were grown in selective liquid media (50 μg/mL ampicillin) overnight, miniprepped, and sequence verified. The resulting plasmid was named pRS315-MmPylRS. pRS315-MmPylRS was then double restriction enzyme digested with SphI and PstI that flank the tRNA_CUA_^Tyr^ gene and SNR52 promoter. A region of vector backbone between the SphI recognition site and the tRNA gene was amplified from pRS315-MmPylRS in addition to the tRNA^ScArg^-tRNA^MmPyl^ region of plasmid SMH-108 (gifted by the Chin Lab at the UK Medical Research Council Laboratory of Molecular Biology). Both PCR-amplified regions were inserted in the digested pRS315-MmPylRS vector via Gibson Assembly. Gibson Assembly reactions were similarly transformed into *E. coli* and grown on selective solid media.

Individual colonies were inoculated into liquid media, miniprepped, and sent for sequence verification. A plasmid containing the correctly assembled insert was named pRS315-MmPylRS-MmtRNA. Cloning for MbPylRS^*20*^ proceeded similarly: MbPylRS and tRNA^ScArg^-tRNA^MmPyl^ were amplified via PCR from plasmids SMH-99 and SMH-108 (both gifted from the Chin Lab at the UK Medical Research Council Laboratory of Molecular Biology). A double digest of pRS315-AcFRS was performed with SacI and PstI. MbPylRS and tRNA^ScArg^-tRNA^MmPyl^ were cloned via Gibson Assembly into the digested vector. Following E. coli transformation, growth of individual colonies in selective liquid media, and miniprepping, the resulting construct was sequence verified and named pRS315-MbPylRS-MmtRNA. Chimeric PylRSs chPylRS(IPYE)^*32*, *33*^ and chAcK3RS(IPYE)^*32*, *33*^ were amplified from pTECH-chPylRS(IPYE) and pTECH-chAcK3RS(IPYE) (gifts from Dieter Söll and David Liu; Addgene plasmid numbers 99222 and 104069), respectively, with overlap regions corresponding to pRS315-MmPylRS-MmtRNA double digested with NcoI and NdeI. The amplified genes were inserted into the digested pRS315-MmPylRS-MmtRNA backbone via Gibson Assembly, transformed into E. coli, inoculated into liquid media, miniprepped, sequence verified, and named pRS315-chPylRS(IPYE) and pRS315-chAcK3RS(IPYE).

Plasmid pRS315-KanRmod-MaPylRS, housing the parent MaPylRS and cognate tRNA_CUA_, was cloned by GENEWIZ. Briefly, the tRNA_CUA_^MaPyl^ sequence was taken from Chin and coworkers’ 2018 publication^*24*^ and the MaPylRS sequence was retrieved from UniProt (M9SC49) from Elsässer and coworkers’ 2018 work with *M. alvus* PylRS.^*25*^ The MaPylRS gene was codon optimized using the IDT Codon Optimizer Tool for *S. cerevisiae* and a gene block including the MaPylRS, a short DNA sequence between the MaPylRS, and the tRNA_CUA_^MaPyl^, was ordered. Genewiz synthesized this insert and performed the cloning process of inserting the synthesized fragment between the NcoI and BstZ17I restriction enzyme sites in a pRS315-KanRmod vector. A mutation was made in the second residue of the gene (T2A) encoding MaPylRS to preserve the NcoI site at the 5’ end of the gene. When inserted between the NcoI and BstZ171 sites in pRS315-KanRmod, this allows for expression of MaPylRS under the GPD promoter and the tRNA_CUA_^MaPyl^ under the SNR52 promoter. Genewiz provided sequencing data verifying the correct assembly of the MaPylRS/tRNA_CUA_^MaPyl^ gene block into the pRS315-KanRmod vector. MaPylRS mutants 1, 2, and 3 were cloned by amplifying the parent MaPylRS from pRS315-KanRmod-MaPylRS with primers containing mutations at specific sites and primers containing 30bp overlap with the digested pRS315-KanRmod vector (SI Tables 1 and 2). pRS315-KanRmod-MaPylRS was digested with restriction enzymes NcoI and NdeI. The PCR fragments and digested vector were analyzed on a DNA gel, extracted, and purified. The fragments were then recombined using Gibson Assembly. Gibson Assembly reactions proceeded for 1 h at 50 °C and then the entire reaction was transformed into chemically competent *E. coli* DH5αZ1. Cells were plated on LB with 34 μg/mL kanamycin and grown at 37 °C overnight. Isolated colonies were inoculated in LB supplemented with kanamycin, grown to saturation, and miniprepped. Isolated plasmids were sequenced via Sanger sequencing. Sequence-verified plasmids were named pRS315-KanRmod-MaPylRS-Mut1, pRS315-KanRmod-MaPylRS-Mut2, and pRS315-KanRmod-MaPylRS-Mut3.

### Yeast transformations, propagation, and induction

pCTCON2-BXG or pCTCON2-BYG and pRS315 plasmids were co-transformed into Zymo-competent *S. cerevisiae* RJY100, plated on selective SD-SCAA –TRP –LEU –URA media, and grown at 30 °C for 3 days. Secretion plasmid transformations proceeded similarly, with pCHA-Donkey1.1-H54TAG-TAA in place of pCTCON2-BXG. For RRE and MMF experiments, three distinct transformants (biological replicates) were inoculated for each plasmid combination. For secretion of Donkey1.1 and Donkey1.1-H54TAG, only one transformant was inoculated in liquid media. Colonies were inoculated in 5 mL SD-SCAA –TRP –LEU –URA supplemented with penicillin-streptomycin (at 100 IU and 100 μg/mL, respectively). All growth and induction liquid cultures were supplemented with penicillin-streptomycin to prevent bacterial contamination. For RRE/MMF experiments, 5 mL cultures were grown at 30 °C with shaking for 2 days and either stored at 4 °C for later use or diluted immediately to OD_600_ = 1 in 5 mL SD-SCAA –TRP –LEU –URA and grown for an additional 4–8 h (OD_600_ = 2–5) before being induced. Inductions were performed in 2 mL volumes of SG-SCAA –TRP –LEU –URA at OD_600_ = 1. Each biological replicate was induced in the absence of ncAAs and in the presence of 10 mM ncAA. Induced cultures were incubated at 20 °C with shaking for 16 h prior to preparation for flow cytometry.

### Flow cytometry data collection and analysis

Following the 16 h induction, 2 MM cells were removed from each culture tube to 96-well V-bottom plates and pelleted. Supernatant was decanted and cells were resuspended in 200 μL PBSA. Two more cycles of centrifugation, decanting, and resuspension were performed for a total of three washes in PBSA. Cells were stored on ice or at 4 °C until resuspension in 200 μL PBSA prior to being analyzed on a flow cytometer. All flow cytometry was performed on an Attune NxT flow cytometer (Life Technologies) at the Tufts University Science and Technology Center.

Data analysis for flow cytometry experiments was performed using FlowJo and Microsoft Excel. Detailed descriptions of how to calculate RRE and MMF are available in prior work.^*30*, *32*^ For the BXG data in Figure 3, we observed carryover of BYG samples into BXG data. To exclude the carryover from analysis, we performed additional gating as follows: a BYG sample was used to draw a gate around WT cells. On the same BYG plot, a gate that would encompass any BFP+ cell except the WT population was drawn (this gate is the “Non-WT” gate, see SI Figure 5 for exact gate positions). The Non-WT gate was applied to all BXG samples and the MFI of BFP and GFP detection from within the Non-WT was used for all BXG samples in Figure 3 when determining RRE and MMF.

For the MFI calculations in Figure 2, the median fluorescence intensity (MFI) of GFP (full-length dual fluorescent reporter) detection was exported from FlowJo for cells displaying above-background levels of BFP detection (i.e., cells exhibiting expression of the reporter) as well as for the population of cells exhibiting only background levels of fluorescence. For each population, the autofluorescence determined from uninduced cells was subtracted and then biological triplicate data was averaged; error bars represent the standard deviation of the biological replicates.

### Protein expression and purification

Transformations and propagations were performed as described above in “Yeast transformations, propagation, and induction.” For Donkey 1.1-H54TAG, 10 colonies were inoculated separately into 5 mL SD-SCAA –TRP –LEU –URA cultures. Cells were grown at 30 °C with shaking for 2–3 days and each 5 mL culture was then diluted into 45 mL SD-SCAA –TRP –LEU –URA and grown an additional 24 h at 30 °C with shaking. All ten 50 mL cultures were pelleted at 2400 rcf for 10 min and then re-suspended separately in 100 mL YPG media with 0.1% BSA supplemented with 10 mM BocK. Induced cultures were incubated at 20 °C with shaking for 4 days.

For Donkey1.1, a single transformant was inoculated in a 5 mL SD-SCAA –TRP –URA culture and grown at 30 °C with shaking for 2 days. The 5 mL culture was diluted in 45 mL SD-SCAA – TRP –URA and grown at 30 °C with shaking for 24 h. 25 mL of the growing culture was used to inoculate a 250 mL culture, which was grown at 30 °C with shaking for an additional 24 h. The 250 mL culture was pelleted at 2400 rcf for 10 min and supernatant was decanted. Cells were resuspended in 500 mL YPG without ncAAs. The induced culture was incubated at 20 °C with shaking for 4 days.

Following induction, cultures were pelleted at 3214 rcf for 30 min. The supernatant was decanted and filtered along with 10X PBS (1.37 M NaCl, 27 mM KCl, 100 mM Na2HPO4, and 18 mM KH2PO4 at pH 7.4) for a 1X final concentration of PBS. Bio-Rad protein purification columns and reservoirs were rinsed with 1X PBS, and 2 mL Protein A slurry (1 mL resin, GenScript) was deposited in the columns. Resin was washed with 1X PBS prior to addition of supernatant. Supernatant was passed over the resin twice. Loaded resin was washed three times with 1X PBS and then protein was eluted in 100 mM glycine, pH 3.0, into a tube with 0.7 mL 1 M Tris pH 8.5. Elution fractions were immediately buffer exchanged using 15 mL 30 kDA molecular weight cutoff devices (Millipore Sigma) into chilled sterile water or 1X PBS using a centrifuge at 4 °C. Proteins were either diluted with 100% glycerol to final 50% v/v glycerol, flash frozen in liquid nitrogen, and stored at –80 °C; or were used immediately for tryptic digests and SDS-PAGE analysis.

### SDS-PAGE

Purified proteins were combined with sterile water, 4X Bolt™ LDS Sample Buffer (Thermo Fisher Scientific), and 10X NuPAGE™ Sample Reducing Agent (Thermo Fisher Scientific). Samples were boiled at 100 °C for 5 min and then loaded into a Bolt™ 4–12%, Bis-Tris, 1.0 mm, 15-well Mini Protein Gel with a Precision Plus Protein™ All Blue Prestained Protein Standard (Bio-Rad) and run for 16 min at 200 V. Gels were washed three times in DI water for 10 min at 60 rpm on an orbital shaker at room temperature, then stained in SimplyBlue™ SafeStain (Thermo Fisher Scientific) for 1 h at 60 rpm on an orbital shaker at room temperature. The gel was de-stained in DI water overnight and imaged the next day on an Azure c400 gel imager (Azure Biosystems).

### Tryptic digests and cleanup for mass spectrometry

−80 °C glycerol stocks of secreted proteins were thawed in a water bath and proteins were buffer exchanged using 0.5 mL 30 kDA molecular weight cutoff devices from Millipore Sigma at 4 °C into sterile water. For proteins used immediately after purification and buffer exchange, the previous step was skipped. 10 μg of each protein was then boiled in PCR tubes at 100 °C for 5 min and allowed to cool to room temperature before 1 μg mass spectrometry-grade trypsin (Trypsin Gold from Promega) was added to each sample. Samples were incubated in a heat block at 37 °C overnight and then cleaned up using C18 ZipTips from Millipore Sigma into 0.1% trifluoroacetic acid/50% acetonitrile. Purified peptide fragments were flash frozen and sent on dry ice to the Koch Institute Biopolymers and Proteomics Core for MALDI mass spectrometry.

## Abbreviations

aaRS: aminoacyl-tRNA synthetase
AcK: *N*_ε_-acetyl-L-lysine
AcrK: acryloyl-L-lysine
BFP: blue fluorescent protein
BocK: Boc-L-lysine
BXG: blue fluorescent protein fused to green fluorescent protein by a flexible linker containing a TAG codon
BYG: blue fluorescent protein fused to green fluorescent protein by a flexible linker
CbzK: *N*_ε_-benzyloxycarbonyl-L-lysine
GFP: green fluorescent protein
LysAlk: 2-Amino-6-(prop-2-ynoxycarbonylamino)hexanoic acid
LysN3: (*S*)-2-amino-6-((2-azidoethoxy)carbonylamino)hexanoic acid
LysZ: *N*_ε_-benzoyl-L-lysine
MALDI: matrix-assisted laser desorption ionization
MaPylRS: *Methanomethylophilus alvus* pyrrolysyl-tRNA synthetase
MbPylRS: *Methanosarcina barkeri* pyrrolysyl-tRNA synthetase
MmPylRS: *Methanosarcina mazei* pyrrolysyl-tRNA synthetase
MMF: maximum misincorporation frequency
ncAA: noncanonical amino acid
NmH2: 3-methyl-L-histidine
OTS: orthogonal translation system
PhK: (*S*)-2-Amino-6-((2-(3-methyl-3H-diazirin-3-yl)ethoxy)carbonylamino)hexanoic acid
PylRS: pyrrolysyl-tRNA synthetase
RRE: relative readthrough efficiency
tRNA: transfer RNA

## Author Information

James A. Van Deventer, Chemical and Biological Engineering Department, 4 Colby Street, Room 267B, Tufts University, Medford, MA 02155, Phone: 617-627-6339

## Acknowledgements

This research was supported by a grant from the the National Institute of General Medical Sciences of the National Institutes of Health (1R35GM133471), and by Tufts University startup funds (to J.A.V.). J.T.S. was supported in part by a National Science Foundation Graduate Research Fellowship (ID: 2016231237). The content of this work is solely the responsibility of the authors and does not necessarily represent the official views of the National Institutes of Health, the National Science Foundation, or Tufts University. The authors would also like to thank Richard Cook, Heather Amoroso, and Alla Leshinsky at the Koch Institute Biopolymers and Proteomics Core for their assistance with mass spectrometry data collection. Additionally, the authors thank A.B. Fitzsimmons for his assistance with mass spectrometry data formatting.

