## Supplementary Information for "Exploration of *Methanomethylophilus alvus* pyrrolysyl-tRNA synthetase activity in yeast"

**Additional sequence information**

**tRNA$_{CUA}^{MaPyl}$ Sequence**
GGGGGACGGTCCGGCGACCAGCGGGTCTCTAAAACCTAGCATAGCGGGGTTCGACACCC CGGTCTCTCG

**MaPylRS Amino Acid Sequence (UniProt M9SC49)**
MTVKYTDAQIQRLREYGNGTYEQKVFEDLASRDAAFSKEMSVASTDNEKKIKGMIANPSRHGL
TQLMNDIADALVAEGFIEVRTPIFISKDALARMTITEDKPLFKQVFWIDEKRALRPMLAPNLYSVM
RDLRDHTDGPVKIFEMGSCFRKESHSGMHLEEFTMLNLVDMGPRGDATEVLKNYISVVMKAA
GLPDYDLVQEESDVYKETIDVEINGQEVCSAAVGPHYLDAAHDVHEPWSGAGFGLERLLTIRE
KYSTVKKGGASISYLNGAKIN

**MaPylRS Codon-optimized DNA Sequence**
ATG ACA GTA AAG TAC ACA GAC GCT CAG ATT CAG CGT CTG AGG GAG TAC GGC
AAC GGT ACT TAC GAG CAA AAA GTC TTT GAA GAC TTG GCT TCC AGG GAT GCG GCT
TTC TCA AAA GAA ATG TCT GTA GCG TCC ACC GAT AAC GAA AAG AAA ATC AAG GGG
ATG ATC GCG AAT CCA AGT AGG CAC GGT CTG ACA CAG TTG ATG AAC GAT ATC GCC
GAT GCA TTG GTT GCA GAG GGC TTC ATT GAG GTT AGA ACG CCA ATC TTC ATA TCT
AAA GAT GCT CTG GCC AGA ATG ACC ATA ACC GAA GAT AAG CCA CTT TTC AAG CAA
GTG TTT TGG ATT GAC GAA AAG CGT GCG TTG AGA CCA ATG TTA GCT CCG AAC CTA
TAT TCT GTG ATG AGA GAC TTG AGA GAT CAC ACC GAC GGA CCA GTC AAG ATT TTT
GAG ATG GGG AGC TGT TTC AGA AAG GAA TCA CAC TCT GGA ATG CAT TTA GAA GAG
TTT ACC ATG CTT AAT CTA GTG GAC ATG GGT CCC AGA GGG GAT GCA ACG GAA GTT
CTA AAG AAC TAC ATC TCT GTA GTG ATG AAA GCT GCT GGC TTG CCC GAC TAC GAT
CTG GTG CAG GAA GAG TCC GAT GTT TAT AAA GAG ACA ATC GAC GTG GAA ATC AAC
GGG CAA GAG GTT TGT TCA GCG GCC GTA GGT CCT CAT TAC TTG GAC GCT GCG
CAT GAT GTT CAC GAA CCT TGG AGC GGA GCA GGT TTT GGC CTA GAG AGG CTG
CTG ACT ATA CGT GAG AAA TAT TCA ACG GTC AAG AAG GGC GGA GCG AGT ATC TCC
TAC CTA AAT GGA GCG AAA ATT AAC

**MaPylRS Codon-optimized DNA Sequence with NcoI mutation at 2nd Position**
ATG **GCA** GTA AAG TAC ACA GAC GCT CAG ATT CAG CGT CTG AGG GAG TAC GGC
AAC GGT ACT TAC GAG CAA AAA GTC TTT GAA GAC TTG GCT TCC AGG GAT GCG GCT
TTC TCA AAA GAA ATG TCT GTA GCG TCC ACC GAT AAC GAA AAG AAA ATC AAG GGG
ATG ATC GCG AAT CCA AGT AGG CAC GGT CTG ACA CAG TTG ATG AAC GAT ATC GCC
GAT GCA TTG GTT GCA GAG GGC TTC ATT GAG GTT AGA ACG CCA ATC TTC ATA TCT
AAA GAT GCT CTG GCC AGA ATG ACC ATA ACC GAA GAT AAG CCA CTT TTC AAG CAA

GTG TTT TGG ATT GAC GAA AAG CGT GCG TTG AGA CCA ATG TTA GCT CCG AAC CTA TAT TCT GTG ATG AGA GAC TTG AGA GAT CAC ACC GAC GGA CCA GTC AAG ATT TTT GAG ATG GGG AGC TGT TTC AGA AAG GAA TCA CAC TCT GGA ATG CAT TTA GAA GAG TTT ACC ATG CTT AAT CTA GTG GAC ATG GGT CCC AGA GGG GAT GCA ACG GAA GTT CTA AAG AAC TAC ATC TCT GTA GTG ATG AAA GCT GCT GGC TTG CCC GAC TAC GAT CTG GTG CAG GAA GAG TCC GAT GTT TAT AAA GAG ACA ATC GAC GTG GAA ATC AAC GGG CAA GAG GTT TGT TCA GCG GCC GTA GGT CCT CAT TAC TTG GAC GCT GCG CAT GAT GTT CAC GAA CCT TGG AGC GGA GCA GGT TTT GGC CTA GAG AGG CTG CTG ACT ATA CGT GAG AAA TAT TCA ACG GTC AAG AAG GGC GGA GCG AGT ATC TCC TAC CTA AAT GGA GCG AAA ATT AAC

**MaPylRS Amino Acid Sequence with NcoI mutation at 2$^{nd}$ Position**
MAVKYTDAQIQRLREYGNGTYEQKVFEDLASRDAAFSKEMSVASTDNEKKIKGMIANPSRHGL
TQLMNDIADALVAEGFIEVRTPIFISKDALARMTITEDKPLFKQVFWIDEKRALRPMLAPNLYSVM
RDLRDHTDGPVKIFEMGSCFRKESHSGMHLEEFTMLNLVDMGPRGDATEVLKNYISVVMKAA
GLPDYDLVQEESDVYKETIDVEINGQEVCSAAVGPHYLDAAHDVHEPWSGAGFGLERLLTIRE
KYSTVKKGGASISYLNGAKIN

**Supplementary figures**

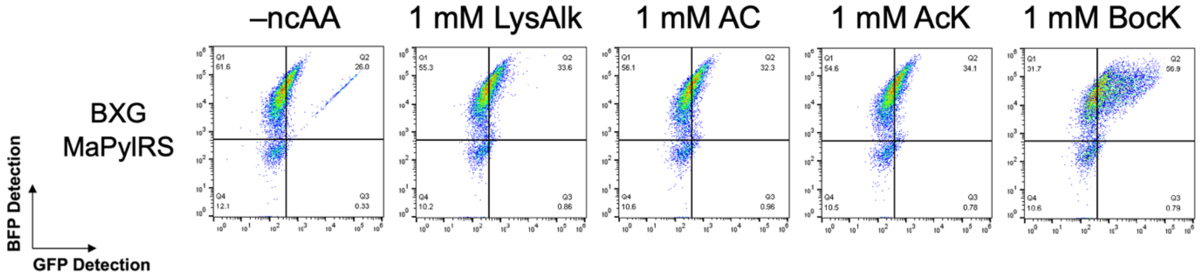


SI Figure 1. Flow cytometry dot plots of wild-type MaPylRS induced in the presence of 1 mM indicated ncAAs with BXG reporter. Experiment was not done with replicates.

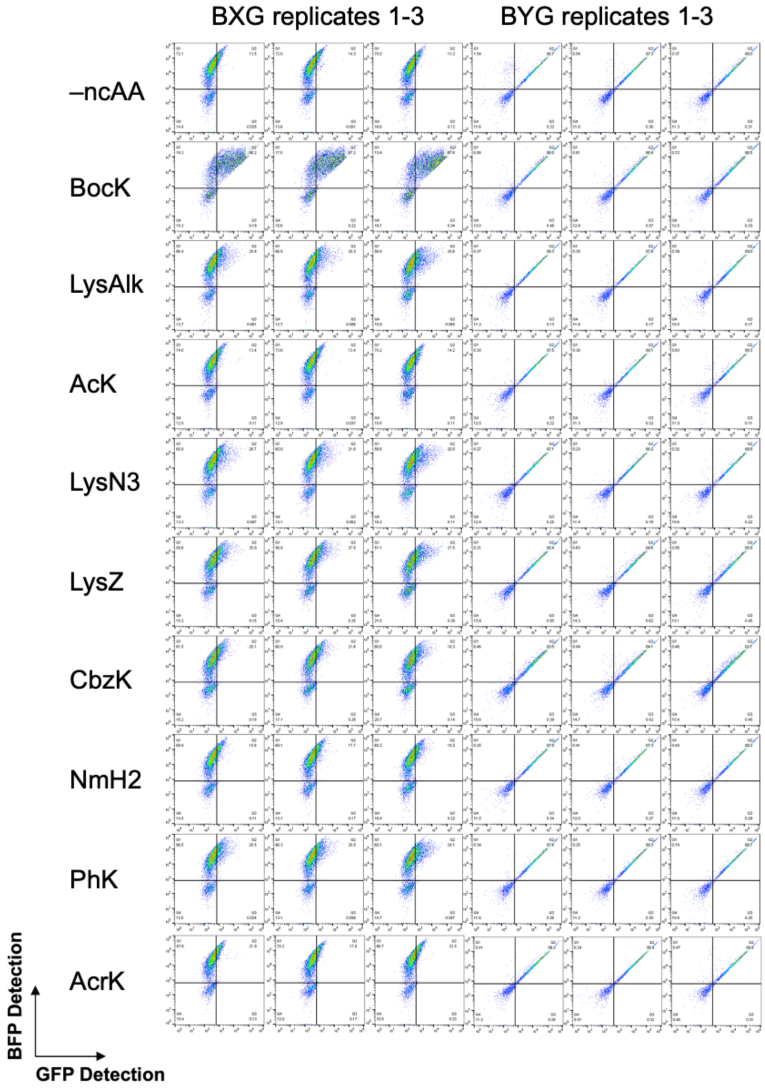


SI Figure 2. Flow cytometry dot plots for data corresponding to RRE and MMF values in Figure 1 of the main text. Biological replicates were used to calculate RRE/MMF and associated error.

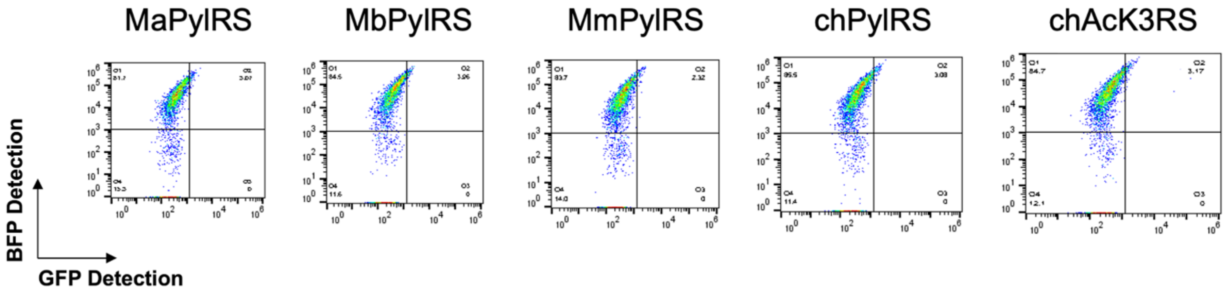


SI Figure 3. Flow cytometry dot plots of negative controls corresponding to data in Figure 2 of the main text. A sample with BXG and each of the PylRSs that was induced in the absence of ncAAs serves as an indicator of how much background cAA misincorporation is expected for each PylRS.

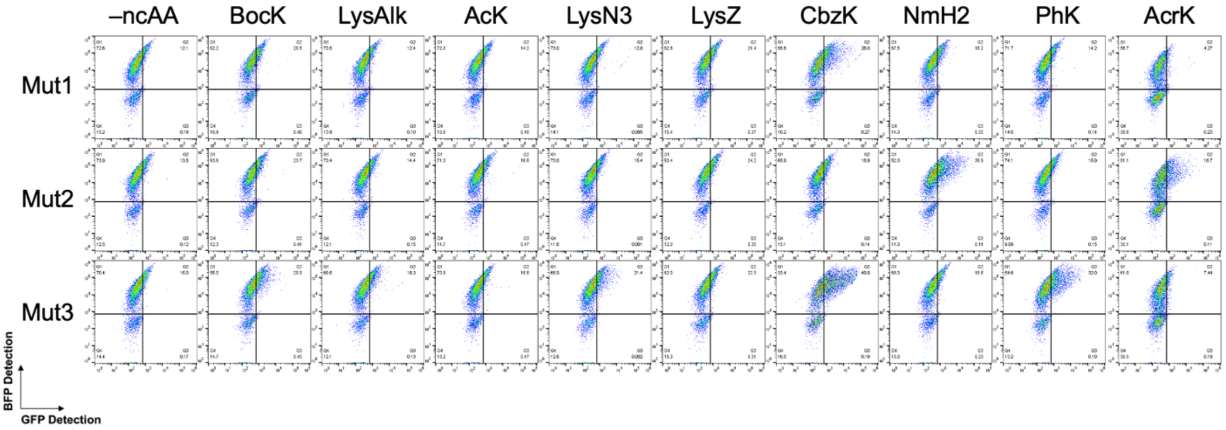


SI Figure 4. Initial flow cytometry results for cells transformed with plasmids encoding constitutively expressed mutant MaPylRSs 1–3 induced in the presence of 10 mM ncAA as indicated.

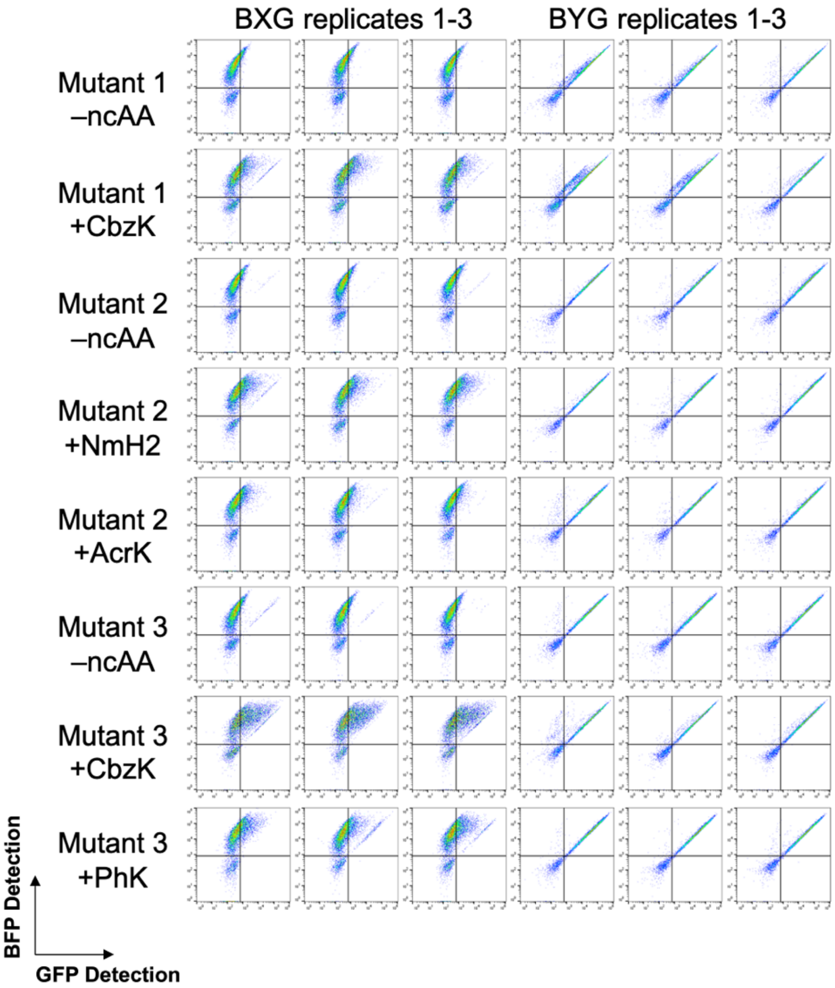


SI Figure 5. Flow cytometry dot plots for data corresponding to RRE and MMF values in Figure 3 of the main text. Biological replicates were used to calculate RRE/MMF and associated error.

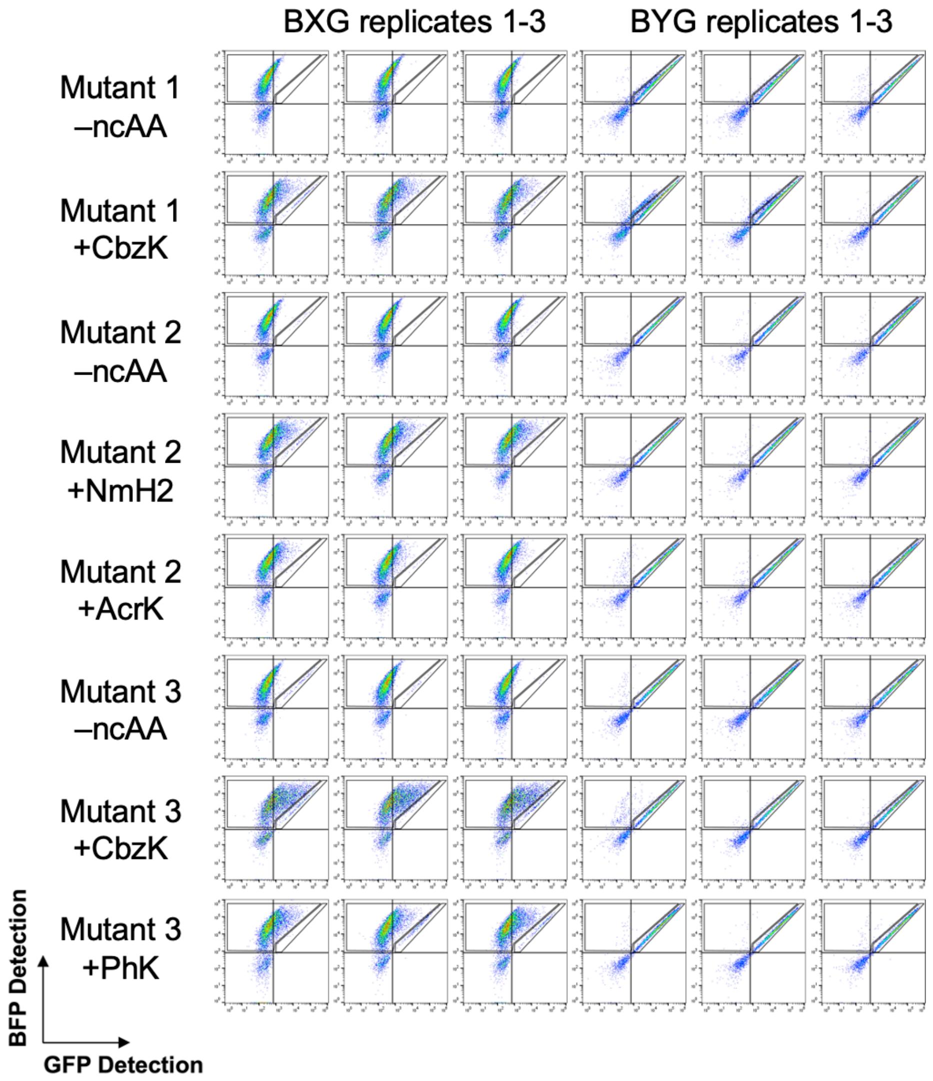


SI Figure 6. Flow cytometry dot plots for data corresponding to RRE and MMF values in Figure 3 of the main text. Due to sample carryover in the autosampler of the flow cytometer, some BYG cells were transferred to BXG samples and needed to be excluded from RRE/MMF calculations so as to avoid skewing the results. Gates used to exclude these carryover events from analysis are presented on these plots.

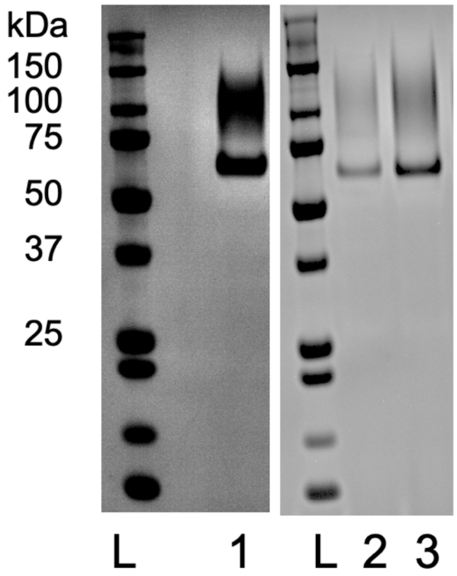


SI Figure 7. SDS-PAGE analysis of proteins post-purification and post-buffer exchange. L is a pre-stained protein ladder; 1 is the wild-type Donkey1.1 reporter; 2 and 3 are the Donkey1.1-H54TAG reporter with MaPylRS induced in the presence of 10 mM BocK with 1 µg and 2 µg masses of protein loaded, respectively. The expected size of the Donkey1.1 reporter is 55.9 kDa.

**Supplementary tables**

Table 1. MaPylRS mutants, mutations, and original source.

| Mutant | NcAA(s) | Mutation(s) | Reference |
|---|---|---|---|
| 1: *Ma*PylRS -MutRS1 | CbzK | Y126M<br>M129G<br>V168T | Willis, J. C. W. and J. W. Chin (2018). "Mutually orthogonal pyrrolysyl-tRNA synthetase/tRNA pairs." Nat Chem 10(8): 831-837. |
| 2: *Ma*PylRS (mut) | NmH2 | L121M<br>L125I<br>Y126F<br>M129A<br>V168F | Beranek, V., et al. (2019). "An Evolved Methanomethylophilus alvus Pyrrolysyl-tRNA Synthetase/tRNA Pair Is Highly Active and Orthogonal in Mammalian Cells." Biochemistry 58(5): 387-390. |
| 3: *Mx1201* $PylRS^{Y126A}$ | TCO*K<br>CpK<br>BCNK<br>AbK | Y126A | Meineke, B., et al. (2018). "Methanomethylophilus alvus Mx1201 Provides Basis for Mutual Orthogonal Pyrrolysyl tRNA/Aminoacyl-tRNA Synthetase Pairs in Mammalian Cells." ACS Chem Biol 13(11): 3087-3096. |

Table 2. Primer names and sequences for MaPylRS mutant cloning. Mutations from WT MaPylRS are noted in pink.

| Mutant | Name | Sequence |
|---|---|---|
| 1 | Mut1-126to129-Fwd | TTG AGA CCA ATG TTA GCT CCG AAC CTA ATG TCT GTG GGT AGA GAC TTG AGA GAT CAC ACC GAC GGA |
| 1 | Mut1-126to129-Rev | TCC GTC GGT GTG ATC TCT CAA GTC TCT TCC CAC AGA CAT TAG GTT CGG AGC TAA CAT TGG TCT CAA |
| 1 | Mut1-168-Fwd | CAT TTA GAA GAG TTT ACC ATG CTT AAT CTA ACG GAC ATG GGT CCC AGA GGG GAT GCA ACG |
| 1 | Mut1-168-Rev | CGT TGC ATC CCC TCT GGG ACC CAT GTC CGT TAG ATT AAG CAT GGT AAA CTC TTC TAA ATG |
| 2 | Mut2-121to129-Fwd | GAC GAA AAG CGT GCG TTG AGA CCA ATG ATG GCT CCG AAC ATT TTT TCT GTG GCT AGA GAC TTG AGA GAT CAC ACC GAC GGA |
| 2 | Mut2-121to129-Rev | TCC GTC GGT GTG ATC TCT CAA GTC TCT AGC CAC AGA AAA AAT GTT CGG AGC CAT CAT TGG TCT CAA CGC ACG CTT TTC GTC |
| 2 | Mut2-168-Fwd | CAT TTA GAA GAG TTT ACC ATG CTT AAT CTA TTT GAC ATG GGT CCC AGA GGG GAT GCA ACG |
| 2 | Mut2-168-Rev | CGT TGC ATC CCC TCT GGG ACC CAT GTC AAA TAG ATT AAG CAT GGT AAA CTC TTC TAA ATG |
| 3 | Mut3-126-Fwd | GCG TTG AGA CCA ATG TTA GCT CCG AAC CTA GCT TCT GTG ATG AGA GAC TTG AGA GAT CAC |
| 3 | Mut3-126-Rev | GTG ATC TCT CAA GTC TCT CAT CAC AGA AGC TAG GTT CGG AGC TAA CAT TGG TCT CAA CGC |
| 1, 2, 3 | MaPylRS_Fwd | CGACGGATTCTAGAACTAGTATGGAGATTT |
| 1, 2, 3 | MaPylRS_Rev | GTGGGGGGAGGGCGTGAATGTAAGCGTGAC |
